# Disruption of Iron Metabolism Resulting from Dmt1/Slc11A2 Deficiency Compromises Notch Protein Degradation and Transcriptional Activation

**DOI:** 10.1101/2025.08.21.671472

**Authors:** Rui Zhang, Somaieh Ahmadian, Jolanda Piepers, Florian Bock, Tom Keulers, Marc A. Vooijs

## Abstract

Notch receptor activation requires γ-secretase-mediated release of Notch intracellular domain 1(NICD1) to regulate gene transcription. Here, we identify the proton-driven Solute carrier 11A2 (Slc11A2) or divalent metal transport protein Dmt1 as an inhibitor of Notch signaling via regulating iron homeostasis and lysosomal integrity. Dmt1 loss reduces ferritin levels and increases labile Fe^2+^, causing elevated reactive oxygen species (ROS) and lipid peroxidation. These changes compromise lysosomal function and impair degradation of S3-Val1744 cleaved NICD1, resulting in its accumulation. Dmt1 has isoforms with or without an iron response element (IRE): Re-expressing Dmt1+IRE robustly increases ferritin heavy-chain (FTH), whereas Dmt1-IRE moderately elevates FTH and ferritin light-chain (FTL), with co-expression further enhancing FTL levels. Restoration of Dmt1 expression rescues ferritin levels, lysosomal activity, and NICD1 degradation while reducing oxidative stress and lipid peroxidation. Notably, Dmt1 deficiency decreases NICD1 binding to RBP-Jκ/CSL and its recruitment to Notch target gene promoters *Hes1* and *Hey1*. Collectively, our findings demonstrate that Dmt1 regulates lysosomal function through iron homeostasis and that lysosomal dysfunction from Dmt1 loss impairs NICD1 degradation and disrupts Notch signaling, linking cellular iron metabolism and Notch pathway activity.

## Introduction

Notch signal transduction is essential for development and cell fate in adult mammalian tissues. Activation begins with proteolytic processing of the receptor: S1-cleavage produces a heterodimeric receptor in the Golgi; Upon ligand binding, S2 cleavage by ADAM protease releases the extracellular domain, followed by S3 cleavage by the γ-secretase complex after which Notch intracellular domain(NICD1)is released into the cytoplasm[1]. NICD1 then translocates to the nucleus, where it associates with the transcription factor RBP-Jκ /CSL and the co-activator MAML to activate Notch target genes.

Endocytosis of the Notch receptor in signal-receiving cells can exert both positive and negative effects on Notch signaling [2]. Endosomal and lysosomal trafficking of both full-length and cleaved Notch receptors is essential for Notch activation [3, 4]. This process facilitates localization of Presenilin1 and 2 [5] and enables pH-dependent γ-secretase cleavage [6], which together regulate NICD1 stability and transcriptional activity [7]. Notably, elevated endosomal and lysosomal pH inhibits Notch signal transduction [8, 9].

Disruption of lysosomal function by lysosomotropic agents, such as chloroquine and bafilomycin A1, has been shown to inhibit Notch-dependent signaling downstream of γ-secretase-mediated cleavage[10, 11]. This indicates that γ-secretase cleavage alone is not sufficient to drive transcriptional activation and highlights the essential role of endolysosomal trafficking in Notch signal transduction.

Previously, we identified the iron transporter Dmt1 as a novel and essential regulator of Notch signaling demonstrating that Dmt1 isoforms serve as binary switches that control Notch dependent cell fate decisions in both normal and tumor cells [11, 12].

Dmt1 (Slc11a2, Nramp2) encodes an iron transporter that mediates non-heme iron uptake in most cell types [13-15]. Its expression is tightly regulated by cellular iron levels through both translational and degradation pathways that involve iron-response elements (IRE) in the 3’ UTR of Dmt1 [13, 16]. Iron homeostasis is regulated by iron regulatory proteins (IRP1 and IRP2) via the iron-responsive element (IRE) pathway. IRPs bind to IRE-containing RNA secondary structures within the untranslated region (UTR) of target mRNA, thereby modulating their translation. These include, among others, Dmt1, transferrin receptor (TFR), and the intracellular iron-binding protein ferritin (FTH) [17, 18]. When iron levels are sufficient, iron binds to IRPs, resulting in dissociation from the IRE, promoting the translation of the target mRNA. However, under iron conditions, IRP binding to IREs inhibits translation of target mRNA[19].

Dmt1 is expressed as four different isoforms; isoforms A and B arise from alternative tissue-specific promoters, while isoforms I and II are with IRE (+) and without IRE (-), respectively [20, 21]. Dmt1 A isoforms are predominantly expressed in absorptive cells of the duodenum and kidney, whereas Dmt1 B isoforms are expressed ubiquitously[19, 20]. Subcellular localization also differs among isoforms: Dmt1 A-I is primarily localized to the plasma membrane, while Dmt1 A-II and Dmt1 B-II are found in recycling endosomes, and Dmt1 B-I is located at the late endosome/lysosome, respectively [21, 22]. Given the differences in expression patterns, subcellular localization of Dmt1 isoforms and effects on Notch signaling, we further investigated how Dmt1 and iron transport influence Notch dependent cell fate decisions.

In this study, we showed that Dmt1-IRE expression enhances ligand-dependent Notch signaling, while Dmt1+IRE inhibits ligand-dependent Notch activation [11]. Silencing of Dmt1 leads to increased lipid peroxidation and lysosomal damage. Moreover, in Dmt1-deficent cells, we observed that intranuclear NICD1 loses its transcriptional activity due to impaired binding to the transcription factor RBP-Jκ.

## Results

### Dmt1 loss results in increased cellular iron levels and increased ROS

We previously established that deletion of Dmt1 in knockout (KO) mouse embryonic fibroblasts (MEF) impaired physiological Notch signaling, which correlated with increased Fe^2+^ and ROS [11]. Using Ferro-orange, a fluorescent dye that binds free intracellular iron, we quantified steady-state iron levels in Dmt1 wildtype (WT) MEF and Dmt1 KO MEFs. In agreement with previous reports, we found that intracellular iron levels were elevated in Dmt1 KO MEF. Consistent with this, pharmacological inhibition of Dmt1 using the small molecule Dmt1 Blocker1 (BL1) reproduced the same phenotype in Dmt1 WT MEF (Figure. 1A).

**Figure. 1.**
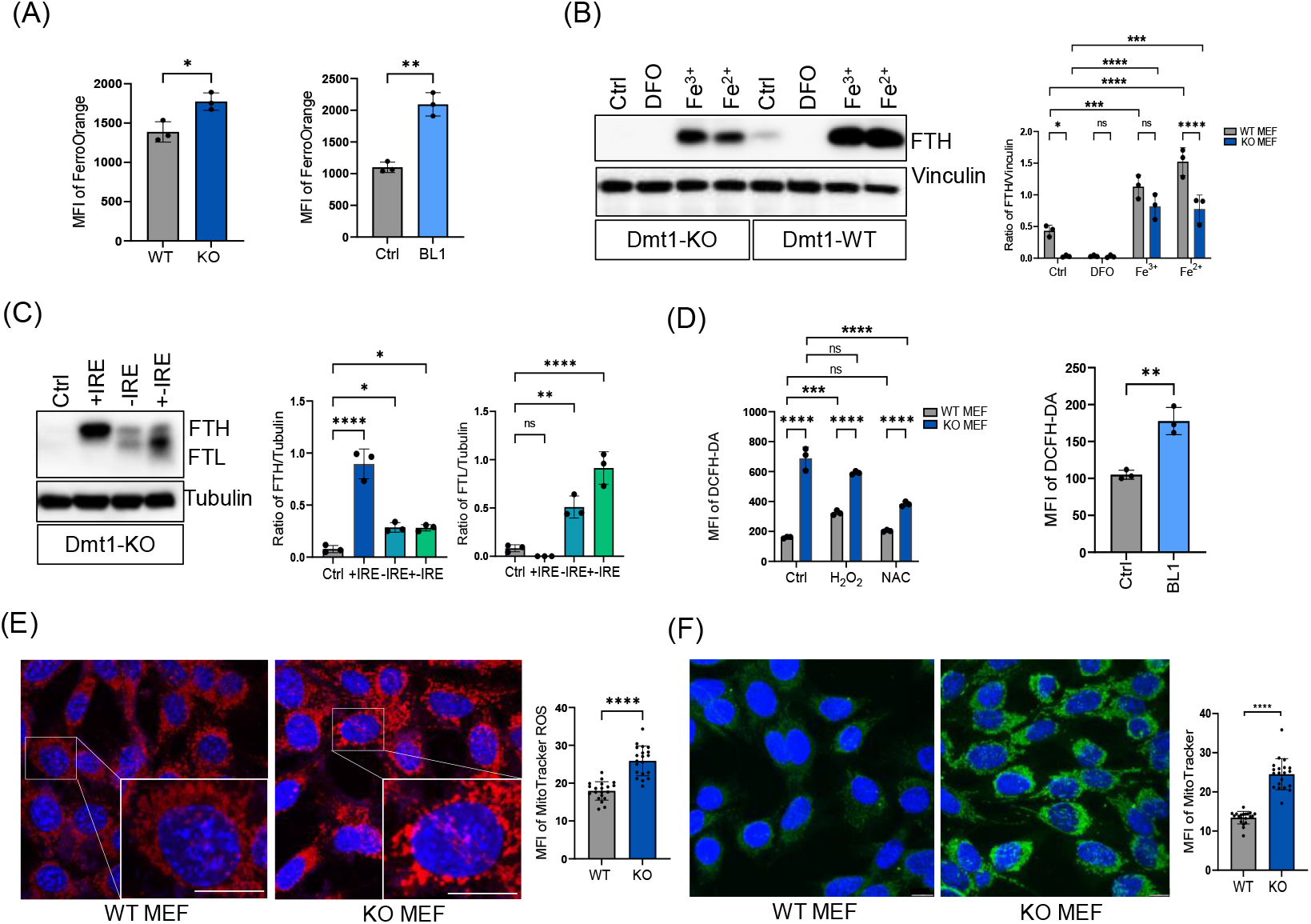
Dmt1 loss results in increased cellular iron levels and increased ROS. **A)** Quantification of intracellular Fe^2+^ levels using FerroOrange fluorescence in Dmt1 WT and KO MEFs, as well as Dmt1 WT MEFs treated by DMSO and Dmt1 inhibitor (BL1). Statistical analysis was performed using unpaired 2-tailed t-test. **B)** Expression of ferritin heavy chain (FTH) in cells treated with DMSO, DFO, Fe^3+^and Fe^2+^, determined by western blot. Statistical analysis was conducted using Two-way ANONA with Šídák’s multiple comparisons test. **C)** FTH and ferritin light chain (FTL) expression in Dmt1 KO MEFs reconstituted with various Dmt1 isoforms, analyzed by western blot. Statistical significance was assessed using one-way ANOVA with Tukey post hoc tests. **D)** ROS generation in Dmt1 WT and KO MEFs treated with DMSO, H_2_O_2_ and NAC and in Dmt1 WT MEFs treated with DMSO and Dmt1 inhibitor (BL1). ROS were determined by DCFHDA detection by flow cytometry. Two-way ANONA with Šídák’s multiple comparisons test and unpaired 2-tailed t-test have been performed respectively. **E)** Representative images and quantification of mitochondria ROS levels in Dmt1 WT and KO MEFs. Quantification was performed on 20 cells from independent experiments. Unpaired 2-tailed t-test has been performed. Scale bar: 10μm. **F)** Representative images and quantification of MitoTracker staining in Dmt1 WT and KO MEFs. Quantification was performed on 20 cells from independent experiments. Scale bar: 10μm. Unpaired 2-tailed t-test has been performed. Data are represented as mean ± standard deviation. Individual dots represent independent observations. Statistical significance \**p*-value<0.05, \*\**p*-value<0.01, \*\*\**p*-value<0.001 and \*\*\*\**p*-value<0.0001.

To assess the impact of disrupted iron homeostasis in Dmt1 KO cells, we examined the expression of the cytosolic iron storage protein. Ferritin is composed of heavy (FTH)- and light-chain (FTL), and its expression is tightly regulated by cytoplasmic iron availability. Dmt1 KO cells exhibited a marked reduction of FTH (Figure. 1B). Supplementing cells with exogenous ferrous Sulfate (Fe^2+^) and ferric sulfate (Fe^3+^) increased FTH levels in both WT and KO cells. As a control, incubation with the iron chelator deferoxamine (DFO) reduced ferritin expression. Importantly, re-expression of Dmt1 isoforms fully restored FTH and FTL expression in Dmt1 KO cells (Figure. 1C, S1A). Notably, re-expression of Dmt1+IRE isoform only could rescue FTH expression, whereas re-expression of Dmt1-IRE rescued both FTH and FTL levels. These results show that Dm1 KO cells exhibit increased total iron levels. However, the observed decrease in ferritin expression suggests reduced availability of iron in the cytoplasm. This implies that iron may be sequestered in other cellular compartments, such as mitochondria or lysosomes, rather than being freely available in the cytosol.

Free intracellular iron pools (Fe^2+^) can generate ROS in the Fenton reaction. We therefore hypothesized that loss of Dmt1 leads to increased intracellular ROS levels. Using dichloro-dihydro-fluorescein diacetate (DCFH-DA) staining of living cells, we detected an increase in steady-state ROS levels in Dmt1 KO cells, which can be rescued by re-expression of Dmt1 (Figure. 1D, S1B). Similarly, intracellular ROS levels also increased in WT MEFs treated with the Dmt1 inhibitor BL1 for 24 hours (Figure. 1D). As Dmt1 is also situated in mitochondria, the primary source of cellular ROS, we determined mitochondrial ROS and mitochondrial membrane potential, using MitoTracker ROS and MitoTracker, respectively. We found that Dmt1 loss increased mitochondrial ROS and mitochondrial membrane potential (Figure. 1E, F).

The level of ROS from mitochondria was modulated by NAC and H_2_O_2_, especially in Dmt1 KO MEFs (Figure. S1C).

### ROS increases induce lipid peroxidation, associated lysosomes

Using electron microscopy, we previously reported that intracellular vesicles, including lysosomes in Dmt1 KO MEFs, exhibited abnormal morphology characterized by ruptured membranes and signs of permeabilization[11]. We hypothesized that this lysosomal damage may be caused by accumulated Fe^2+^ and increased ROS, resulting in lipid peroxidation of the lysosomal membrane. Using fluorescent dye BODIPY^TM^ 581/591 C11, we observed that lipid peroxidation was increased in Dmt1 KO MEFs. This effect was further intensified by H_2_O_2_ treatment, markedly enhancing lipid peroxidation (Figure. 2A), similar to the response observed in wildtype MEF treated with the Dmt1 inhibitor BL1 (Figure. 2B). Furthermore, re-expression of Dmt1+-IRE decreased lipid peroxidation (Figure. S2A). Fluorescent imaging revealed that lipid peroxidation was specifically increased in the lysosomes of Dmt1 KO MEFs, indicating localized oxidative stress (Figure. 2C). This was accompanied by a rise in LAMP1-positive foci, suggesting an increase in lysosome number. Consistent with elevated oxidative stress, glutathione peroxidase 4 (GPX4) levels were also increased in Dmt1 KO cells, an effect that was reversed by re-expression of the Dmt1+IRE (Figure. S2B).

**Figure. 2.**
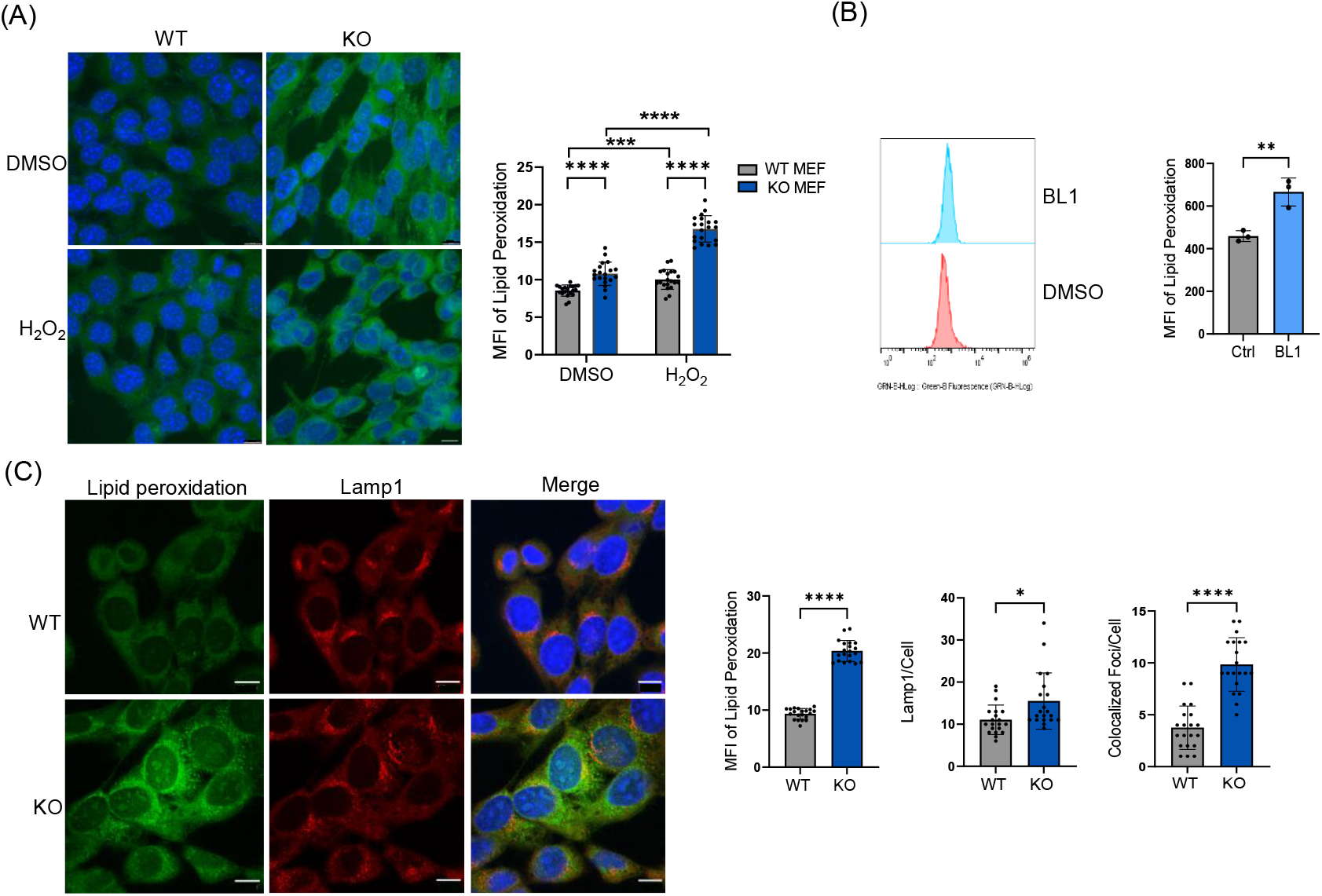
Dmt1 deficiency promotes lipid peroxidation in lysosomes. **A)** Representative images and quantification of lipid peroxidation staining in Dmt1 WT and KO MEFs treated with DMSO and H_2_O_2_. 20 cells from independent experiments were quantified by Image J. Scale bar: 10μm. Two-way ANONA with Šídák’s multiple comparisons test has been used. **B)** Lipid peroxidation (Probe: BODIPY^TM^ 581/591 C11) analyzed by flow cytometry in Dmt1 WT MEFs treated with DMSO and Dmt1 inhibitor (BL1). Statistical significance was determined by unpaired 2-tailed t-test. **C)** Representative images and quantification of lipid peroxidation (BODIPY, green) co-localization with lysosomes (Lamp1, red) in Dmt1 WT and KO MEFs. Co-localization analysis was performed on 20 cells from independent experiments using ImageJ. Scale bar: 10 μm. Statistical analysis was conducted using an unpaired two-tailed t-test. Data are represented as mean ± standard deviation. Individual dots represent independent observations. Statistical significance \**p*-value<0.05, \*\**p*-value<0.01, \*\*\**p*-value<0.001 and \*\*\*\**p*-value<0.0001.

### Loss of Dmt1 impairs lysosomal acidification and lysosomal intracellular activity

To further investigate the integrity of the lysosomal pool, we assessed the presence of intact and functional acidic lysosomes. To assess this, we used LysoTracker Red DND-99, a lipophilic, basic fluorescent dye that selectively labels acidic organelles (pH < 5.0) such as lysosomes in living cells. Live-cell imaging revealed a significant reduction in total integrated LysoTracker signal per cell in Dmt1 KO MEFs (Figure. 3A), consistent with decreased intensity of LysoTracker in Dmt1 KO MEFs via flow cytometry analysis (Figure. S3A) which was rescued by re-expression of Dmt1 (Figure. S3B).

**Figure. 3.**
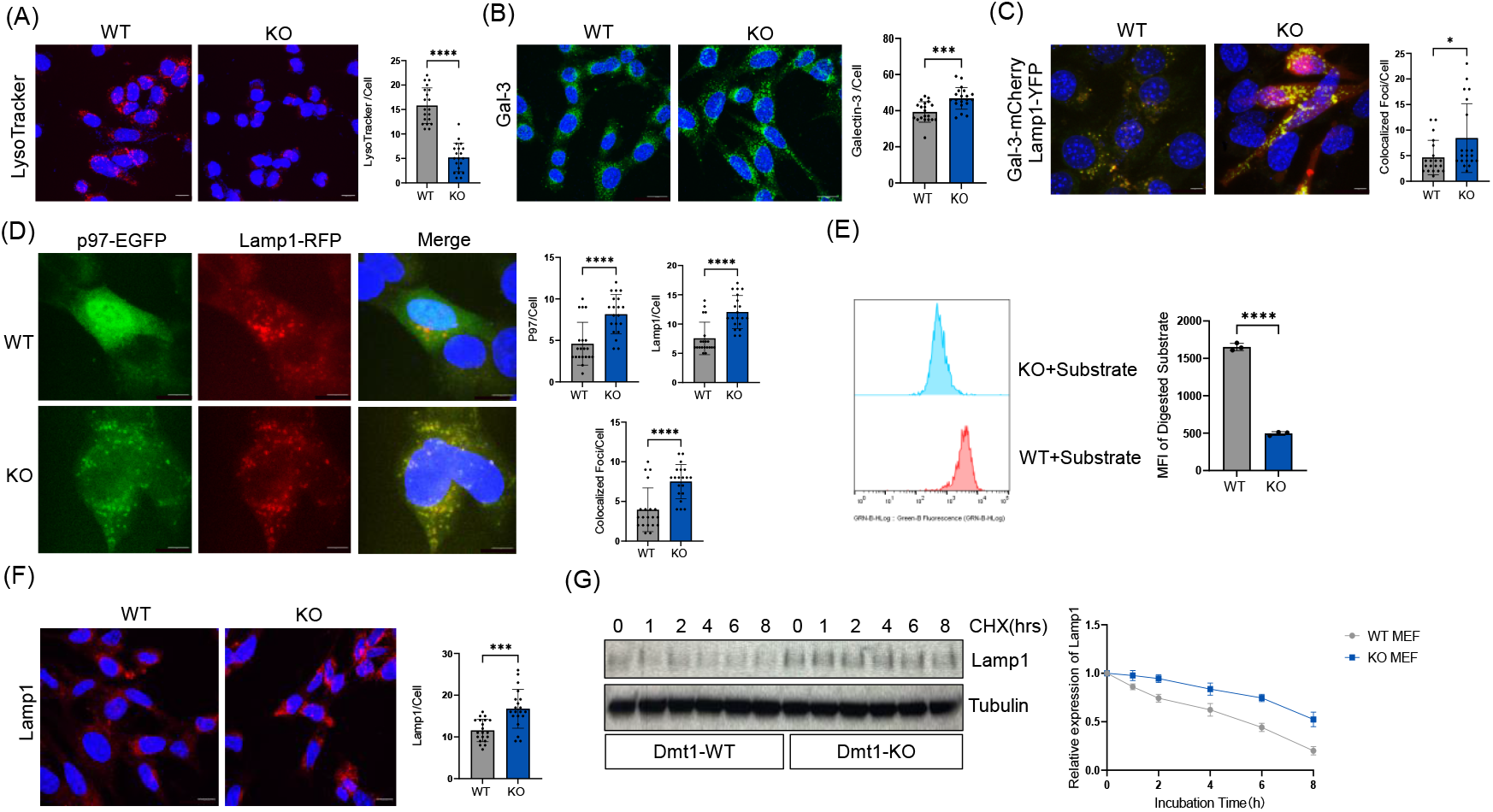
Loss of Dmt1 impairs lysosomal acidification and lysosomal intracellular activity. **A)** Representative images and analysis of LysoTracker intensity in Dmt1 WT and KO MEFs. 20 cells from independent experiments were quantified by Image J. Unpaired 2-tailed t-test has been performed. Scale bar: 10 μM. **B)** Representative images and analysis of Galectin-3 intensity in Dmt1 WT and KO MEFs. 20 cells from independent experiments were quantified. Scale bar: 10μm. Unpaired 2-tailed t-test has been performed. **C)** Representative images and quantification of colocalization between Gal-3-mCherry and Lamp1-YFP in Dmt1 WT and KO MEFs. 20 cells from independent experiments were analyzed. Scale bar: 10μm. Unpaired 2-tailed t-test has been performed. **D)** Representative images and analysis of colocalization between p97-EGFP and Lamp1-RFP in Dmt1 WT and KO MEFs. 20 cells from independent experiments were quantified. Scale bar: 10μm. Unpaired 2-tailed t-test has been performed. **E)** Lysosomal intracellular analysis in Dmt1 WT and KO MEFs by flow cytometry. Unpaired 2-tailed t-test has been performed. **F)** Representative images and quantification of Lamp1 in Dmt1 WT and KO MEFs. 20 cells from independent experiments were quantified. Scale bar: 10μm. Unpaired 2-tailed t-test has been performed. **G)** Lamp1 degradation in Dmt1 WT and KO MEFs treated with DMSO and Cycloheximide (CHX) analyzed by western blot. The samples were collected after 0,2,4,6,8 hours of incubation with CHX and analyzed. Data are represented as mean ± standard deviation. Individual dots represent independent observations. Statistical significance \**p*-value<0.05, \*\**p*-value<0.01, \*\*\**p*-value<0.001 and \*\*\*\**p*-value<0.0001.

Next, we analyzed the recruitment of Galectin-3 (Gal-3), a well-established marker of lysosomal membrane permeabilization (LMP). Endogenous Gal-3 puncta were significantly increased in Dmt1 KO cells (Figure. 3B), and co-localized with Lamp1 in cells co-transfected with Gal-3-mCherry and Lamp1-YFP (Figure. 3C, S3C), confirming lysosomal damage. Cells mitigate the harmful consequences of lysosomal leakage through autophagy-mediated removal of damaged lysosomes. Additionally, p97 translocates to damaged lysosomes and is essential for the removal of damaged lysosomes by autophagy [25]. Consistent with this, we observed a significant increase in the colocalization of p97-GFP and Lamp1-RFP in Dmt1-deficient MEFs (Figure. 3D), further confirming LMP.

Lysosomal acidification is essential for the activity of degradative enzymes like cathepsins, which require an acidic pH for substrate degradation. To assess the impact of Dmt1 deficiency on lysosomal function, we utilized flow cytometry to measure the degradation of a fluorescent lysosomal substrate reporter. Our analysis confirmed a significant impairment in substrate degradation in Dmt1-deficient cells (Figure. 3E) which was rescued by Dmt1 re-expression (Figure. S3D).

Furthermore, we observed an accumulation of the resident lysosomal protein Lamp1 in Dmt1 KO cells (Figure. 3F), suggesting either defective lysosomal degradation or increased protein stability. To differentiate between these, we treated Dmt1 KO cells with cycloheximide (CHX) to inhibit new protein synthesis and monitored Lamp1 turnover following CHX removal using immunoblotting. Dmt1-deficient cells displayed increased state levels of Lamp1 protein but a delayed rate of Lamp1 degradation (Figure. 3G), consistent with impaired lysosomal proteolysis.

Taken together, these data suggest that loss of Dmt1 leads to lysosomal membrane damage caused by increased lipid peroxidation, impaired lysosomal acidification, and compromised lysosomal proteolytic function.

### Dmt1 deficiency directly affects Notch degradation and transcriptional activation

We previously demonstrated that Dmt1-deficient cells exhibit defective Notch1 signaling, characterized by NICD1 stabilization without activation of Notch target genes after physiological ligand stimulation[11]. To determine whether NICD1 degradation is impaired, we examined its turnover. Notch degradation is primarily regulated by CDK8 phosphorylation and FBXW7/Cdc4-mediated ubiquitination and proteasomal degradation [26, 27]. Treatment with the proteasome inhibitor MG132 resulted in a rapid accumulation of NICD1 within 2 hours in WT cells (Figure. 4A). However, in Dmt1 KO MEFs, MG132 did not further stabilize NICD1, suggesting that proteasomal degradation of NICD1 is impaired. To directly assess NICD1 turnover, we performed pulse-chase experiments with CHX, which revealed that NICD1 degradation was significantly delayed in Dmt1 KO cells (Figure. 4B), like the degradation defect observed for Lamp1.

**Figure. 4.**
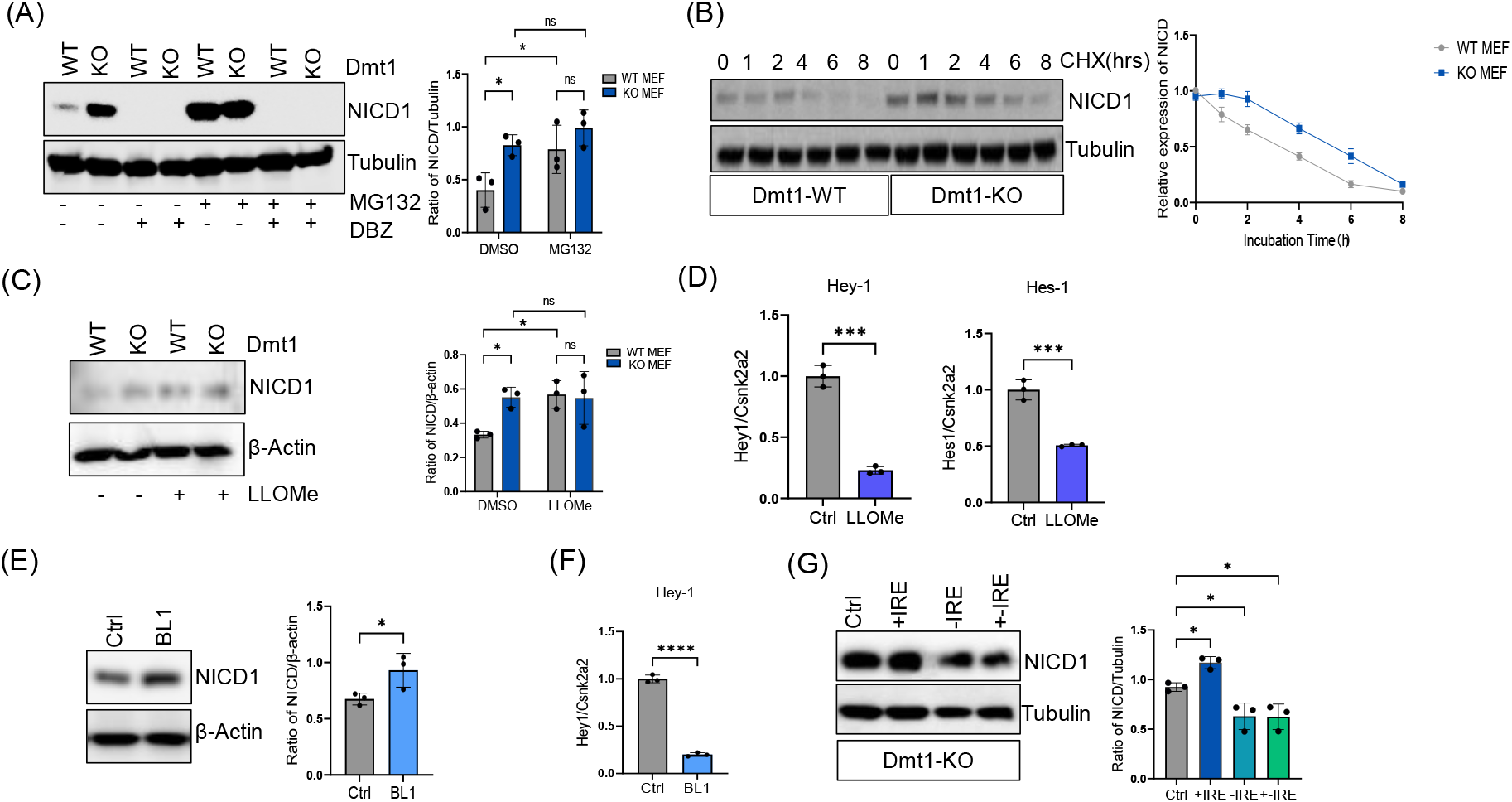
Dmt1 deficiency impairs Notch degradation and transcriptional activation. **A)** NICD1expression in Dmt1 WT and KO MEFs treated with proteasome inhibitor (MG132) and γ-secretase inhibitor (DBZ) analyzed by western blot. Two-way ANONA with Šídák’s multiple comparisons test has been used. **B)** NICD1 degradation in Dmt1 WT and KO MEFs treated by DMSO and Cycloheximide (CHX) analyzed by western blot. The samples were collected after 0,2,4,6,8 hours of incubation with CHX and analyzed. **C)** NICD1 expression in Dmt1 WT MEFs treated by DMSO and Leu-leu methyl ester hydrobromide (LLOMe) analyzed by western blot. Two-way ANONA with Šídák’s multiple comparisons test has been used. **D)**Notch target genes (*Hey1* and *Hes1*) expression in Dmt1 WT MEFs treated with DMSO and LLOMe analyzed by qPCR. Unpaired 2-tailed t-test has been performed. **E)** NICD1 expression in Dmt1 WT MEFs treated with DMSO and Dmt1 inhibitor (BL1) analyzed by western blot. Unpaired 2-tailed t-test has been performed. **F)** Notch target gene (*Hey1*) expression in Dmt1 WT MEFs treated with DMSO and BL1 analyzed by qPCR. Unpaired 2-tailed t-test has been performed. **G)** NICD1 expression in Dmt1 KO MEFs with the re-expression of Dmt1 isoforms analyzed by western blot. One-way ANOVA with Tukey post hoc tests has been performed. Data are represented as mean ±standard deviation. Individual dots represent independent observations. Statistical significance \**p*-value<0.05, \*\**p*-value<0.01, \*\*\**p*-value<0.001 and \*\*\*\**p*-value<0.0001.

To further evaluate whether an intact lysosomal pool is required for Notch signaling, lysosomes were damaged using the lysosome-damaging agent l-leucyl-l-leucine methyl ester (LLOMe). Consistent with our observation in Dmt1 KO cells, lysosomal damage induced by LLOMe resulted in NICD1 stabilization in ligand stimulated wildtype MEF (Figure. 4C) and reduced mRNA expression of the Notch target genes *Hey1* and *Hes*1 in WT MEFs (Figure. 4D). Additionally, pharmacological inhibition of Dmt1 with BL1 induced NICD1 stabilization (Figure. 4E) and *Hey1* mRNA suppression (Figure. 4F). Importantly, re-expression of Dmt1 restored lysosomal substrate cleavage and reinstated NICD1 degradation (Figure. 4G), leading to recovery of Notch pathway activation, as evidenced by *Hey1* mRNA expression (Figure. S4A). Together, these results directly link Dmt1 and lysosomal function to physiological Notch1 signaling.

### Dmt1 deficiency blocks nuclear NICD1-RBP-Jκ /CSL complex formation and decreases recruitment of p62/ SQSTM1 to NICD1

Given the impaired Notch1 signaling observed in Dmt1-deficient cells, we examined the subcellular localization of endogenous Notch1 following physiological ligand stimulation. Immunofluorescent staining for Val144-cleaved NICD1 revealed nuclear accumulation in both WT and Dmt1 KO cells (Figure. 5A). Despite elevated NICD1 levels and its nuclear localization in Dmt1 KO cells, Notch signaling remained suppressed, prompting us to examine defects in the Notch transcriptional complex. To assess the NICD1 chromatin association, subcellular fractionation was performed. In Dmt1 KO cells, NICD1 was predominantly found in the soluble nuclear fraction, with reduced levels in the insoluble chromatin-bound fraction (Figure. 5B), suggesting impaired recruitment to target gene promoters. In the canonical Notch pathway, NICD1 forms a transcriptional complex with the DNA-binding protein RBP-Jκ/CSL and the co-activator Mastermind (MAML) to drive the expression of Notch target genes. RBP-Jκ/CSL protein levels remained unchanged in Dmt1 KO cells; however, immunoprecipitation (IP) revealed a marked reduction in NICD1-RBP-Jκ complex formation (Figure. 5C). To determine whether this defect impaired Notch target gene regulation, we performed chromatin immunoprecipitation (ChIP). Dmt1 KO cells exhibited significantly reduced NICD1 occupancy at the *Hes*1 and *Hey*1 promoters, compared to physiological Notch signaling in WT MEFs (Figure. 5D). Furthermore, we found that the binding of the adapter p62/SQSTM1 to NICD1 was reduced in Dmt1 KO MEFs (Figure. 5E). Taken together, the results provide a direct mechanistic explanation for the reduced assembly of the transcriptional complex and activation of Notch target genes in the absence of Dmt1.

**Figure. 5.**
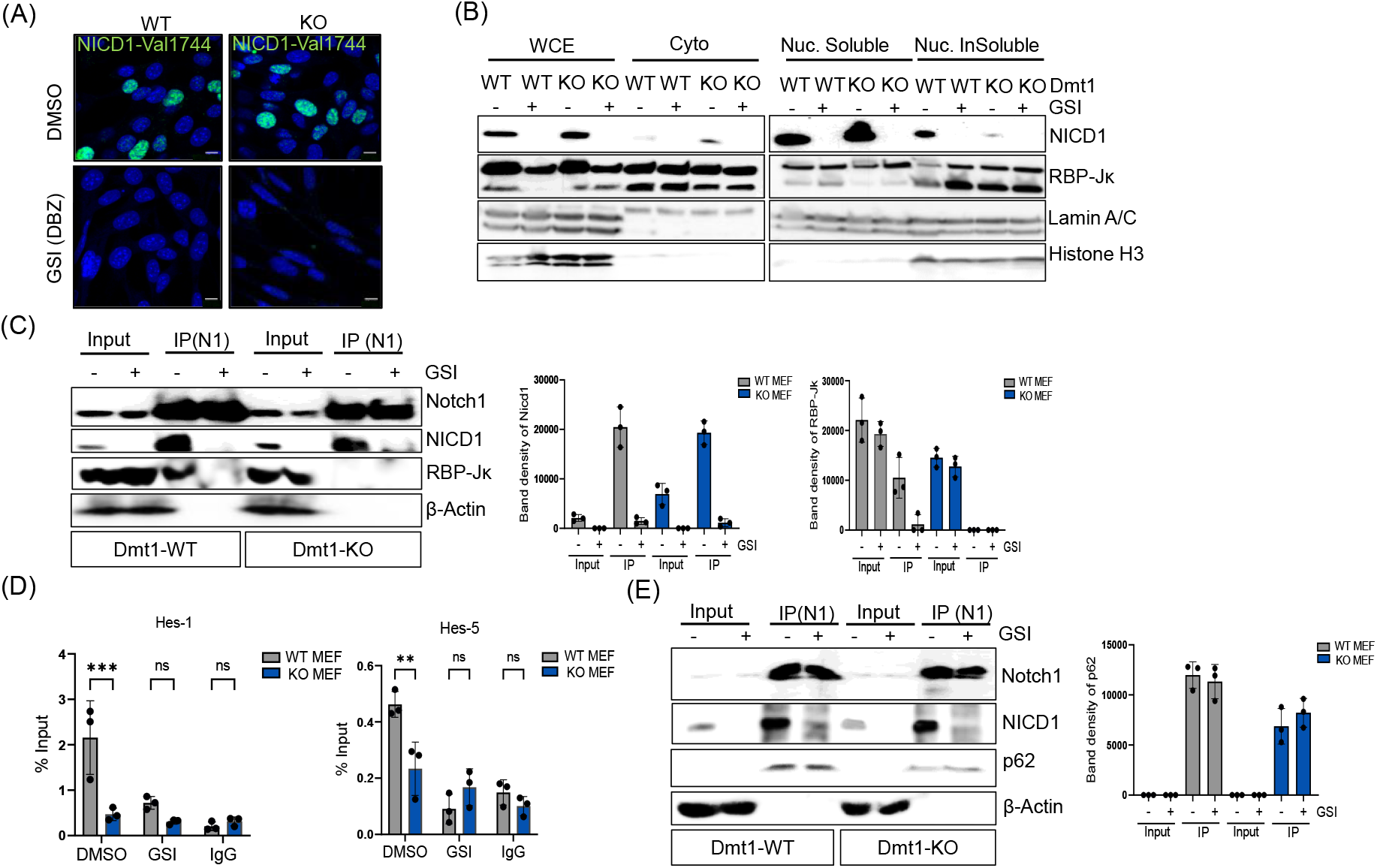
Dmt1 deficiency blocks nuclear NICD1-RBP-Jκ complex formation and decreases recruitment of p62 to NICD1. **A)** Representative images of colocalization including NICD1 and nucleus in Dmt1 WT and KO MEFs treated with DMSO and γ-secretase inhibitor (DBZ). Scale bar: 10μm. **B)** NICD1 and RBP-Jκ localization in different cell components of Dmt1 WT and KO MEFs analyzed by cell fractionation and western blot. WCE=Whole cell extract. Cyto=cytoplasm. Nuc=nucleus. **C)** NICD1-RBP-Jκ complex formation analyzed by immunoprecipitation (IP) and western blot. **D)** Notch target genes (*Hes1*and*Hes5*) regulation in Dmt1 WT and KO MEFs analyzed by chromatin immunoprecipitation (ChIP). Two-way ANONA with Šídák’s multiple comparisons test has been used. **E)** Recruitment of p62 to NICD1 in Dmt1 WT and KO MEFs analyzed by IP and western blot. Data are represented as mean ± standard deviation. Individual dots represent independent observations. Statistical significance \**p*-value<0.05, \*\**p*-value<0.01, \*\*\**p*-value<0.001 and \*\*\*\**p*-value<0.0001.

## Discussion

Our findings establish Dmt1 as a key regulator of iron homeostasis, lysosomal integrity, and physiological Notch signaling, thereby linking endogenous Notch receptor regulation to cellular iron metabolism and proteostasis[28]. Dmt1 loss leads to iron accumulation and elevated ROS, which induces oxidative stress and disrupts lysosomal function through impaired acidification and membrane permeabilization, thereby ultimately compromising proteolytic degradation and disrupting Notch signaling. Here we show that, although available in the nucleus, the Notch intracellular domain fails to assemble with the RBP-Jκ /CSL transcription complex required for target gene transcription, resulting in impaired Notch signaling.

In this study, we demonstrate that silencing Dmt1 leads to impaired Notch signaling, which is associated with disruption of the endolysosomal system. In line with previous reports[11], we show that Dmt1-deficient cells exhibit lysosomal damage, evidenced by a reduction in intact, functional lysosomes, as determined by pH-dependent LysoTracker staining, increased membrane peroxidation, and increased Galectin-3 localization to lysosomes. These findings indicate lysosomal membrane permeabilization, likely triggered by elevated levels of ROS caused by disrupted iron metabolism.

Dmt1 is an essential transmembrane iron transporter. Its isoforms containing or lacking an iron-responsive element (+IRE and –IRE, respectively) are localized to the plasma membrane, late endosomes/lysosomes, and recycling endosomes, where they contribute to cellular iron homeostasis. Silencing Dmt1 expression leads to an increase in total cellular Fe^2+^ levels, as detected by FerroOrange, accompanied by decreased levels of ferritin, the main iron storage protein. Since ferritin mRNA contains an iron-responsive element (IRE), its translation typically increases in response to elevated cytoplasmic iron. However, the paradoxical combination of elevated free Fe^2+^ and reduced ferritin in our study suggests impaired iron storage. Given that Dmt1 is essential in exporting Fe^2+^ from lysosomes to the cytoplasm, we hypothesize that Fe^2+^ accumulates in the lumen of lysosomes of Dmt1-deficient cells and a loss of ferritin in lysosomes, contributing to lysosomal dysfunction and oxidative stress. Further studies are needed to investigate subcellular differences of Fe^2+^ in WT and KO MEFs.

We hypothesize that loss of Dmt1 causes lysosomal iron overload, resulting in lysosomal damage and oxidative stress through the Fenton Reaction[29, 30]. In this reaction, Fe^2+^ catalyzes the conversion of hydrogen peroxide into highly reactive radicals which can damage cellular content such as nucleic acids, proteins, and membranes. Notably, the increased expression of glutathione peroxidase 4 (GPX4) in KO MEFs may represent a compensatory response to ROS-induced damage. However, the antioxidant capacity in KO MEFs was impaired, as evidenced by a significant increase in lipid peroxidation and mitochondrial ROS following hydrogen peroxide treatment.

Remarkably, in addition to the high oxidative state of Dmt1 KO cells, these cells also are defective in Notch signaling. Previous studies have shown that cells with a disrupted endo-lysosomal system are defective in notch signaling [9] [8] [31], highlighting the dependency of notch signaling on the endolysosomal trafficking and function. Functional studies further demonstrated that inhibition of the lysosomal system reduces notch signaling, in part through defective γ-secretase activity leading to decreased NICD availability in the nucleus[32]. Notably, genome-wide knockout screens performed in our group[11], identified Dmt1 as the most potent regulator of Notch1 signaling, ranking above other lysosome associated proteins. This suggests a specific and prominent role for Dmt1 and iron metabolism in regulating Notch signaling, beyond their general functions in lysosomal homeostasis [33].

Interestingly, in our study, we observed that NICD1 is still transported to the nucleus in Dmt1-deficient cells, yet Notch target gene activation is impaired. This suggests that, beyond receptor cleavage and nuclear translocation, either i) additional post-translational modifications of NICD are required for its transcriptional activity, or ii) iron is directly or indirectly required for the assembly or function of the NICD/RBP-Jκ transcriptional complex.

One known posttranslational modification required for Notch signaling is hydroxylation, a process dependent on iron availability through iron-dependent hydroxylases[34-36]. In line with this, the disruption of iron metabolism in our study coincides with reduced expression of Notch target genes. Therefore, one possible explanation for this finding is that iron-dependent hydroxylases are impaired due to disrupted iron availability, thereby reducing Notch target gene activation.

Autophagy has also been implicated in the regulation of NICD stability. For instance, it has been shown that the autophagy adaptor protein p62 binds to the ubiquitinated PEST domain of NICD1, thereby promoting its degradation via autophagy [37]. Another study elucidated a potential mechanism in which the process of selective autophagic degradation of NICD1 may be initiated in the nucleus through its binding with LC3B.LC3B was shown to co-localize with the lysosomal protein Lamp1, suggesting that NICD is selectively trafficked to autolysosomes for degradation [38]. Our previous study also indicated that NICD1 is present in autophagosomes, as shown by co-staining with LC3B [10], and it has been reported that autophagy regulates Notch degradation to modulate stem cell development [39]. Taken together with our findings, reduced p62 binding to NICD1 may contribute to the inhibition of NICD1 degradation via autophagy or proteasomal degradation.

Additionally, some histone-modifying enzymes, such as certain histone demethylases, are iron-dependent and require iron as a cofactor for their catalytic activity, whereas histone deacetylases (HDACs) typically depend on zinc or other metals. For instance, histone demethylase KDM5A, an iron dependent demethylase and essential part of the NICD/RBP-Jκ repression mechanism[40]. Further research is needed to investigate how iron deficiency in Dmt1 KO cells affects iron-dependent enzymes required for Notch stability and transcriptional activation.

In conclusion, our study demonstrates that Dmt1 regulates Notch signaling through its roles in iron metabolism, lysosomal integrity, and NICD1 modification. These findings reveal a mechanistic link between Dmt1 and Notch pathway activation, establishing a novel connection between iron homeostasis, lysosomal function, and Notch signaling. Given these interactions, we propose that therapeutic strategies targeting iron-related disorders, lysosomal storage diseases, or Notch-associated pathologies should carefully consider potential synergistic effects and unintended consequences arising from this crosstalk.

## Materials and methods

### Compounds

The reagents were used as described in the previous study [11]. Cells were treated with 100μM Iron(II) sulfate heptahydrate(FeSO_4_·7H_2_O(Sigma,cat.#F7002) for 24h, 100 μM Iron(III) sulfate hydrate(Fe_2_(SO_4_)_3_)(Sigma,cat.#307718) for 24h, 100 μM Deferoxamine mesylate salt(Sigma,cat.#D9533)for 24h, 1.3μM Dmt1 blocker 1 (Cymit Quimica,cat.#TM-T11063) for 24h, 125μM Leu-leu methyl ester hydrobromide (LLOMe)(Sigma,cat.#16689-14-8) for 1h, 100μM Hydrogen peroxide(Sigma,cat.#H1009) for 30 min and 50 μg/mL Cycloheximide(Sigma,cat.#C7698).

### Plasmids

pQCXIH-Dmt1b+IRE-HA and pQCXIH-Dmt1b-IRE-Flag were constructed as described previously[11]. The following plasmids were ordered from Addgene: pEGFP-p97(#85670), Lamp1-RFP (#1817), Lamp1-YFP (#1816) and mCherry-Gal3(#85662).

### Cell lines

Jagged2-stimulated mNramp2(Dmt1) WT/KO MEFs were generated as described previously[11]. Dmt1b+IRE, Dmt1b-IRE and Dmt1+-IRE re-expression MEFs (Jagged2-stimulated) were generated by transduction with pQCXIH-Dmt1b+IRE-HA, pQCXIH-Dmt1b-IRE-Flag and both, respectively. Retrovirus was produced by transfecting expression vector together with packaging plasmids(pBS-CMV-gagpol) (Addgene, cat. #35614) and envelop plasmid(pCMV-VSV-G) (Addgene, cat. #8454) into HEK293FT cells. Cell supernatants were harvested 48 hours after transfection. To obtain stable cell lines, Jagged2-stimulated Dmt1 WT/KO MEFs were infected for 24 hours with retroviral supernatants in the presence of 10μg/mL Polybrene (Sigma, cat. #H9268). 48 hours after transfection, Dmt1b+IRE re-expression MEFs were screened with 2mg/mL G418 disulfate salt (Sigma, cat. #G8168), Dmt1b-IRE re-expression MEFs were screened with 1mg/mL Hygromycin B (Gibco^TM^, cat. #10687010), and Dmt1b+-IRE MEFs were screened with 2mg/mL G418 disulfate salt and 1mg/mL Hygromycin B. The surviving cells were pooled and passaged before use.

All cells were maintained in DMEM (Dulbecco’s Modified Eagle’s medium, high glucose) (Gibco) with 10% FBS under cell culture conditions (37°C,5%CO_2_) and regularly tested for mycoplasma contamination.

### Immunofluorescence and image processing

The procedure was conducted according to previously described methods [11]. The information on antibodies is described as follows: Lamp1(Abcam, cat. #ab24170,1:1000) and Galectin-3(Proteintech, cat. #60207-1-Ig). For cells treated with fluorescent dyes, samples after fixation were counterstained with DAPI, mounted with mounting media and visualized directly with confocal microscope.

For Val1744(NICD1) staining, the immunofluorescence was performed using Alexa Fluor^TM^488 Tyramide SuperBoost^TM^ kit, goat anti-rabbit IgG (Invitrogen^TM^, cat. #B40922), based on the manufacturer’s protocol. In short, MEFs were seeded in 8-well µ-Slides (ibidi, cat. #80826) at a density of 2 × 10^4^ cells per well. Cells were fixed with 4% paraformaldehyde for 10 min, permeabilized with 0.1% Triton X-100 for 10 min and quenched with 50 mM glycine in PBS. Endogenous peroxidase activity was blocked using 3% hydrogen peroxide for 60 min at room temperature. Following blocking with 10% goat serum, cells were incubated with rabbit anti-Val1744 antibody (Cell Signaling, cat. #4147S,1:1000) (1:100 in 2% BSA + 10% goat serum in 0.1% Triton X-100) for 1 h at room temperature. After washing, cells were incubated with HRP-conjugated goat anti-rabbit secondary antibody for 1 h, followed by Tyramide signal amplification using Alexa Fluor 488-conjugated Tyramide for 2–10 min. The reaction was stopped with a stop reagent, and cells were counterstained with DAPI. Cells were mounted with fluorescent mounting medium and stored at 4 °C for imaging.

Mitochondria were analyzed by 200nM MitoTracker Green (Cell Signaling, cat. #9074S) incubating 30 min at 37 °C. Mitochondria ROS was measured by 500nM MitoTracker Red CMXRos (Invitrogen^TM^, cat. #M7513) incubating 2 hours. The cell was fixed with 3.7% formaldehyde in complete growth medium at 37°C for 15 min.

### Immunoblotting

The procedure was carried out as previously described [11]. Details of the primary antibodies are as follows: rabbit anti-cleaved Notch1(Val1744, D3B8) (Cell Signaling, cat.#4147S,1:1000), rabbit anti-Lamp1(Abcam, cat.#ab24170,1:1000), rabbit anti-FTH (Cell Signaling, cat.#3998S,1:1000), mouse anti-Vinculin (Sigma, cat.#V9131,1:1000), mouse anti-Tubulin (Sigma, cat.#T7451,1:1000), mouse anti-Flag (Sigma, cat.#F3165,1:1000), rabbit anti-HA (Sigma, cat.#H6908,1:1000), rabbit anti-GPX4(Proteintech,cat.#30388-1-AP), rabbit anti-β-Actin(Proteintech, cat.# 81115-1-RR), rabbit anti-RBP-Jκ (ThermoFisher, cat.#MA5-35438) for cell fractionation, rabbit anti-Lamin A(Sigma, cat.#L1293), rabbit anti-Histone H3(Cell Signaling, cat.#9715), rat anti-Notch1(Cell Signaling, cat.#3447), mouse anti-RBP-Jκ (Santa Cruz, cat.#sc-271128) for IP, and rabbit anti-p62(Cell Signaling, cat.#5114).Band intensity was quantified using ImageJ software.

### Sample processing for flow cytometry

Iron (Fe^2+^) was detected in Dmt1 WT and KO MEFs with iron sensing dye FerroOrange (Cell Signaling, cat. #36104) according to the manufacturer’s protocol. Shortly, cell culture media was removed, and cells were washed with 1×PBS and subsequently incubated with 1 μmol/L FerroOrange working solution for 30 min at 37°C. Next, cells were washed with 1×PBS and analyzed by flow cytometry.

ROS level was analyzed by incubating cells with 5μM 2’,7’-dichlorofluorescin diacetate (DCFHDA) (Sigma, cat. #35845) for 30min at 37°C, and cells were washed with 1×HBSS analyzed by flow cytometry.

Lipid peroxidation was detected by BODIPY^TM^ 581/591 C11(Invitrogen^TM^, cat. #D3861). Cells were treated by 5μM BODIPY^TM^ 581/591 C11 for 30min at 37°C and then washed by 1×PBS. The fluorescence of cells was evaluated at 488nm-excitation and 510nm-emission wavelengths.

LysoTracker intensity was measured by LysoTracker Red DND-99(Invitrogen^TM^, cat. #L7528). Cells were treated by 50nM of LysoTracker Red DND-99 for 1h at 37°C and then washed by 1×PBS. The fluorescent intensity was analyzed with excitation and emission maximum of 577/590 nm.

Lysosomal intracellular activity was measured by Lysosomal Intracellular Activity Assay Kit (Abcam, cat. #ab234622) based on the manufacturer’s protocol. In short, cells media was replaced with 0.5% FBS media and 15 uL substrate was added into 1mL media. After 1h incubation at 37°C, cells were washed twice in 1mL ice-cold 1×Assay Buffer. Finally, cells were analyzed by Guava EasyCyte™ 12 flow cytometry at 488nm excitation wavelength.

### Quantitative RT-PCR

We performed the procedure according to methods described previously [11]. mRNA abundance was measured by the usage of forward and reverse primers (Table.1).

**Table. 1:**
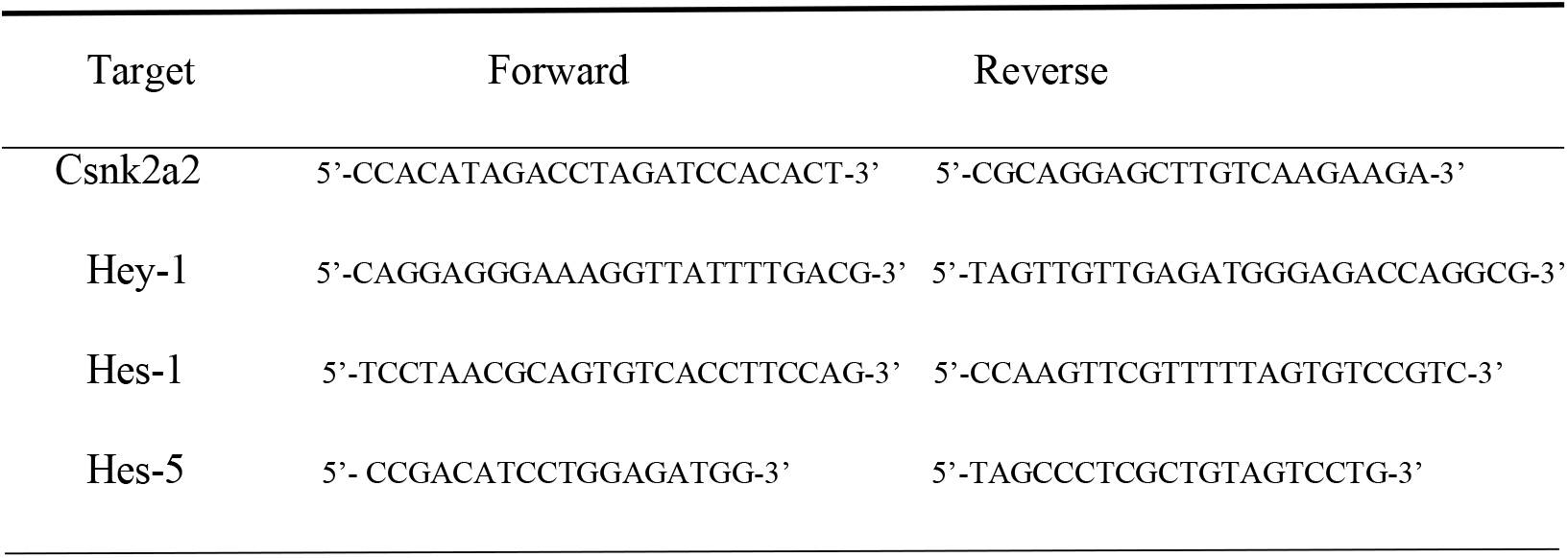
Primers used for qPCR determination.

### Cell fractionation

The fractionation protocol was described previously[23] with some adaptations. For nuclear-cytoplasmic fractionation, MEFs were harvested by trypsinization, resuspended in complete medium to inhibit proteolysis, and centrifuged at 500 × g for 4 min at 4 °C. Cell pellets were washed twice with ice-cold PBS and centrifuged again under the same conditions. To initiate cell swelling, pellets were resuspended in hypotonic buffer (20 mM Tris-HCl pH 7.4, 10 mM KCl, 2 mM MgCl_2_, 1 mM EGTA, 0.5 mM DTT, 0.5 mM PMSF) and incubated on ice for 3 min. NP-40 was then added to a final concentration of 0.1%, and samples were incubated for an additional 3 min on ice. The homogenate was centrifuged at 2000 × g for 5 min at 4 °C to separate the nuclear (pellet) and cytoplasmic (supernatant) fractions. The cytoplasmic supernatant was clarified by centrifugation at 15,000 × g for 3 min at 4 °C to remove residual debris.

The nuclear pellet was washed two times with isotonic buffer (20 mM Tris-HCl pH 7.4, 150 mM KCl, 2 mM MgCl_2_, 1 mM EGTA, 0.5 mM DTT, 0.5 mM PMSF) then once with isotonic buffer supplemented with 0.1% NP-40, incubated on ice for 5–10 min, and centrifuged at 1000 × g for 3 min. For nuclear subfractionation, the washed nuclei were lysed in RIPA buffer (25 mM Tris-HCl pH 7.4, 150 mM NaCl, 0.1% SDS, 0.5% sodium deoxycholate, 0.1% NP-40, protease inhibitors). Samples were incubated on ice for 30 min and then centrifuged at 2000 × g for 3 min at 4 °C to separate the RIPA-soluble nucleosolic fraction (supernatant) from the RIPA insoluble chromatin bound pellet.

For further insoluble nuclear fractionation, pellets were treated with DNase I in DNase buffer (20 mM Tris-HCl pH 7.4, 100 mM NaCl, 4 mM MgCl_2_, 1 mM CaCl_2_, 0.1% NP-40, 0.5 mM DTT, 0.5 mM PMSF) for 30 min on ice and subsequently sonicated to release DNA-associated proteins.

### Immunoprecipitation (IP)

Cells were lysed with ice-cold lysis buffer (Thermo Scientific^TM^, cat. #87787) containing protease inhibitor cocktail (Roche, cat. #11873580001). Cell lysates were centrifuged at 14000g for 15 min at 4°C, and the supernatant was collected for concentration test by Bradford (Bio-Rad, cat. #5000001). Protein was mixed with Notch1 antibody (Cell Signaling, cat. #3608S) at 4 °C overnight. Protein G beads (Invitrogen^TM^, cat. #10003D) were immunoprecipitated with protein and antibody complex at 4 °C 1 h. The beads were washed 3 times with lysis buffer and then heated at 100°C in SDS sample buffer for immunoblotting.

### Chromatin Immunoprecipitation

Sample preparation was performed as described previously[24]. Specifically, the chromatin pull-down antibody and beads were the same as IP. Normal Rabbit IgG antibody (Cell Signaling, cat. #2729S) was used as the control. The sequences of oligonucleotides used as ChIP primers are listed in Table.1.

### Statistical analysis

Data were presented as mean±SD with three biological repeats. Statistical analyses were performed using graphpad prism 9. The unpaired 2-tailed t-test was used for single comparisons, one-way ANOVA with Tukey post hoc tests was used for intergroup comparisons, and two-way ANONA with Šídák’s multiple comparisons test was used for comparison across multiple experimental groups. Results were considered significant if \**p*-value<0.05, \*\**p*-value<0.01, \*\*\**p*-value<0.001 and \*\*\*\**p*-value<0.0001.

## Abbreviations

Dmt1: Divalent metal transporter1
Slc11A2: solute carrier family11, member2
MEF: mouse embryonic fibroblasts
FTH: ferritin heavy-chain
FTL: ferritin light-chain FTL
Lamp1: lysosomal-associated membrane proteins1
ROS: reactive oxygen species
GSI: γ-secretase inhibitor
DBZ: γ-secretase inhibitor
IRE: iron-responsive element
IRP: iron regulatory protein
TFR: transferrin receptor
NICD: Notch intracellular domain
DCFH-DA: dichloro-dihydro-fluorescein diacetate
CHX: cycloheximide
GPX4: glutathione peroxidase4
LLOMe: l-leucyl-l-leucine methyl ester
NAC: n-acetyl cysteine
DFO: deferoxamine
BL1: Dmt1 inhibitor

## Acknowledgement

This research was supported by the Nederlandse Organisatie voor Wetenschappelijk Onderzoek under Grant OCENW. M20.155 and the China Scholarships Council under Grant 202108310006.

## Conflicts of interest

The authors declare no conflict of interest.

## Author contribution

RZ, SA and JP performed the experiments and analysed the experimental data. RZ drafted the manuscript and designed the figures under the supervision of FB, TK and MV. FB, TK and MV contributed to the design and implementation of the research. All authors discussed the results and commented on the manuscript.

## Data availability statement

The data that support the findings of this study are available from the corresponding author upon reasonable request.

---

**Supplementary figure S1.**
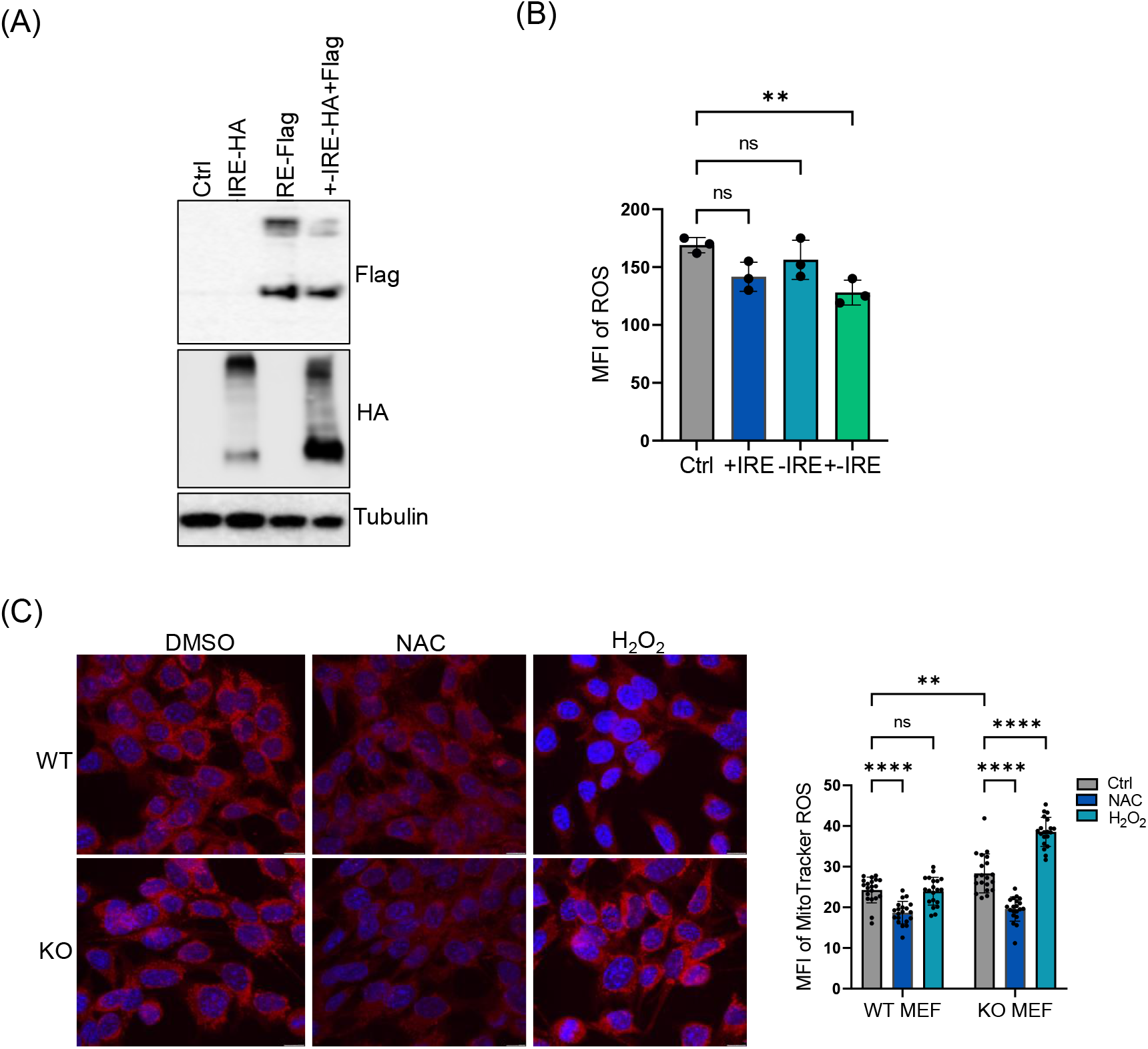
Re-expression of Dmt1 isoforms and ROS analysis. **A)** Re-expression of Dmt1 isoforms in Dmt1 WT MEFs analyzed by western blot. **B)** ROS generation analysis in Dmt1 KO MEFs with the re-expression of Dmt1 isoform via DCFHDA detection by flow cytometry. One-way ANOVA with Tukey post hoc tests has been performed. **C)** Representative images and quantification of mitochondria ROS staining in Dmt1 WT and KO MEFs treated with DMSO, H_2_O_2_ and NAC. 20 cells from independent experiments were quantified by Image J. Scale bar: 10μm. Two-way ANOVA with Šídák’s multiple comparisons test has been used. Data are represented as mean ±standard deviation. Individual dots represent independent observations. Statistical significance \**p*-value<0.05, \*\**p*-value<0.01, \*\*\**p*-value<0.001 and \*\*\*\**p*-value<0.0001.

**Supplementary figure S2.**
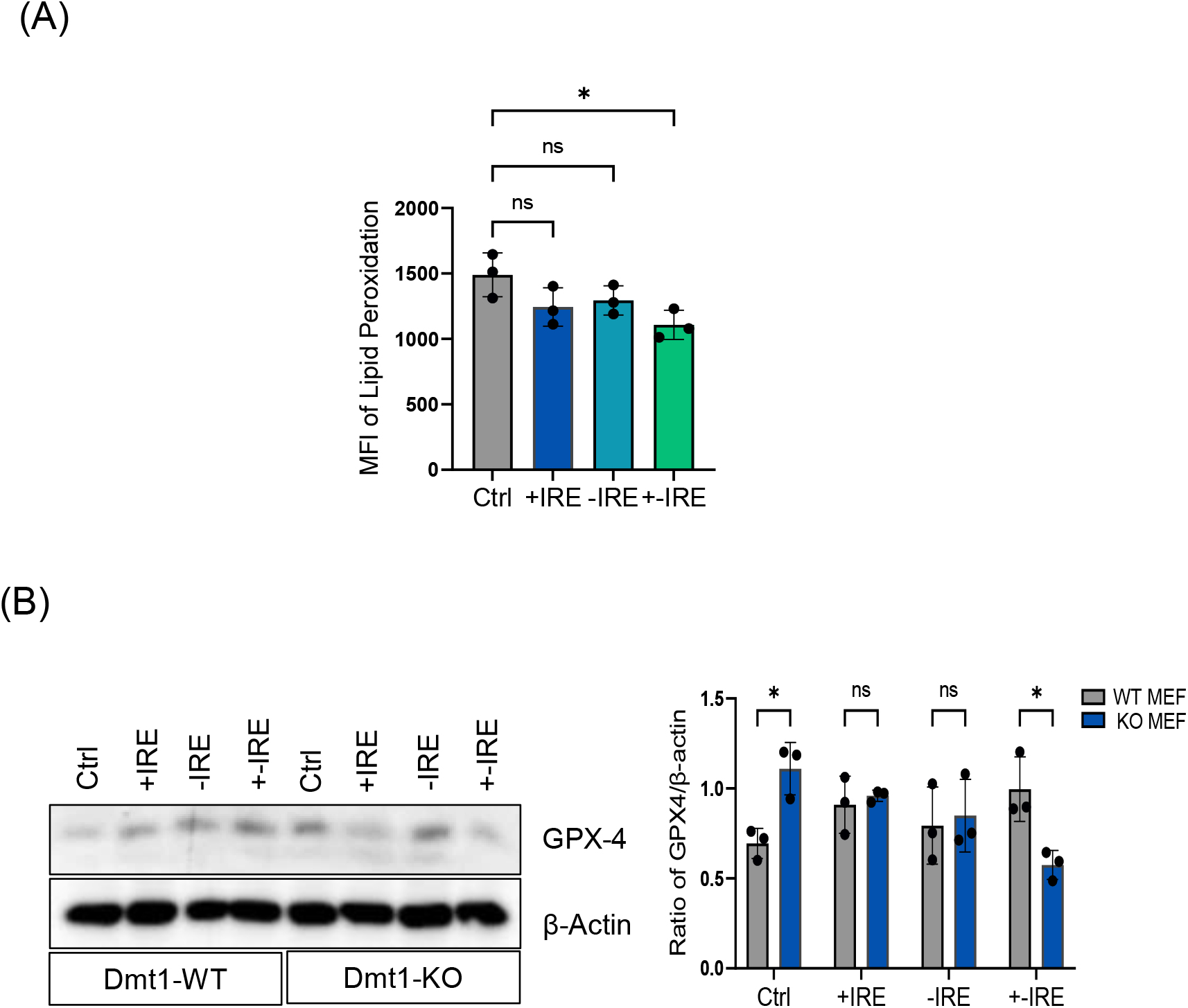
MFI of lipid peroxidation and GPX4 expression in Dmt1 isoforms re-expression cells. **A)** Lipid peroxidation analyzed by flow cytometry in Dmt1 KO MEFs with the re-expression of Dmt1 isoforms. One-way ANOVA with Tukey post hoc tests has been performed. **B)** GPX4 expression in Dmt1 WT and KO MEFs with the re-expression of Dmt1 isoforms analyzed by western blot. Two-way ANONA with Šídák’s multiple comparisons test has been used. Data are represented as mean ± standard deviation. Individual dots represent independent observations. Statistical significance \**p*-value<0.05, \*\**p*-value<0.01, \*\*\**p*-value<0.001 and \*\*\*\**p*-value<0.0001.

**Supplementary figure S3.**
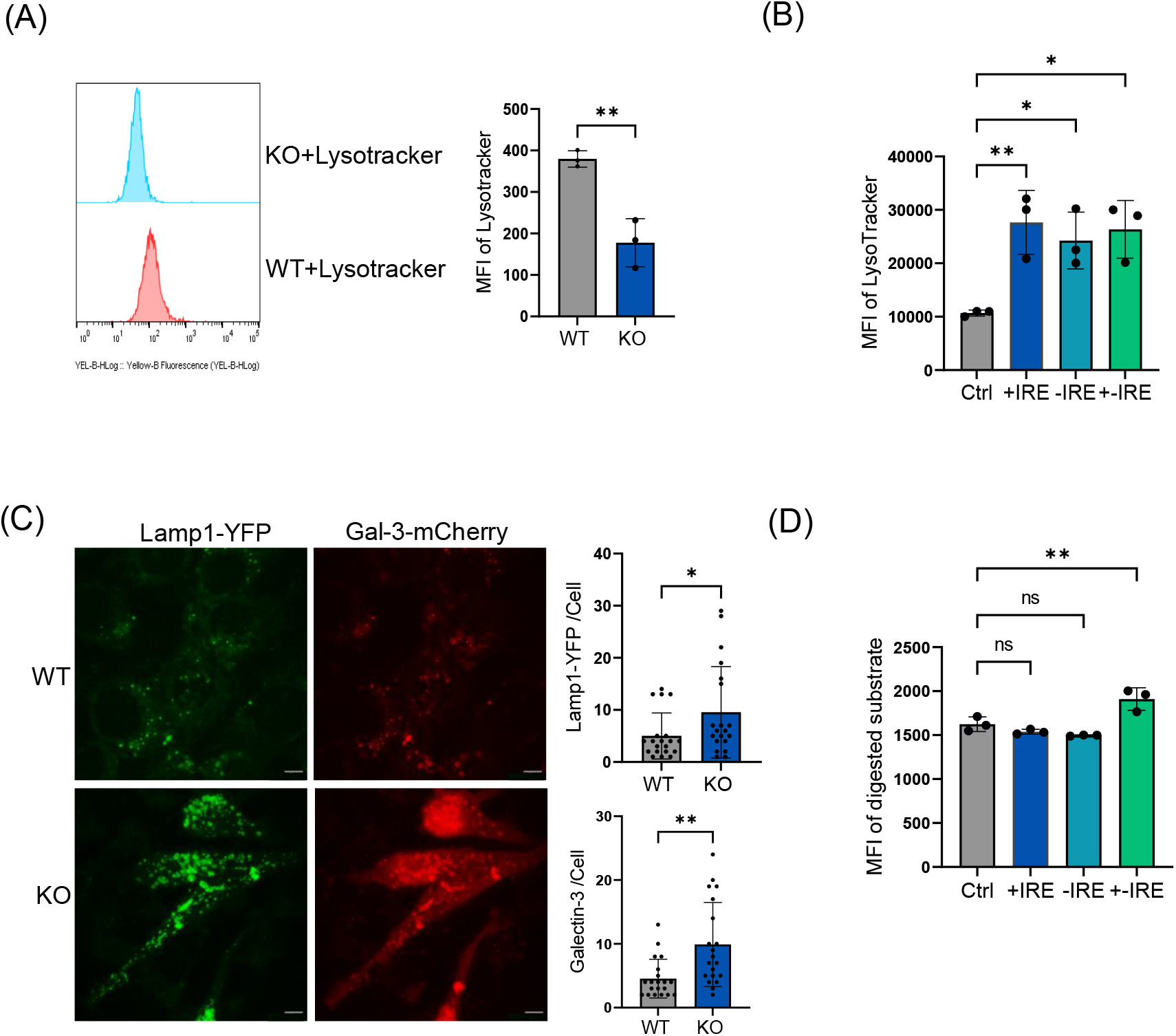
Lysosomal stability and functional analysis. **A)** LysoTracker intensity analysis in Dmt1 WT and KO MEFs by flow cytometry. Unpaired 2-tailed t-test has been performed. **B)** LysoTracker intensity analysis in Dmt1 KO MEFs with the re-expression of Dmt1 isoforms by flow cytometry. One-way ANOVA with Tukey post hoc tests has been performed. **C)** Representative images and analysis of Lamp1-YFP and Gal-3-mCherry in Dmt1 WT and KO MEFs. 20 cells from independent experiments were quantified by Image J. Scale bar: 10μm. Unpaired 2-tailed t-test has been performed. **D)** Lysosomal intracellular analysis in Dmt1 KO MEFs with the re-expression of Dmt1 isoforms by flow cytometry. One-way ANOVA with Tukey post hoc tests has been performed. Data are represented as mean ±standard deviation. Individual dots represent independent observations. Statistical significance \**p*-value<0.05, \*\**p*-value<0.01, \*\*\**p*-value<0.001 and \*\*\*\**p*-value<0.0001.

**Supplementary figure S4.**
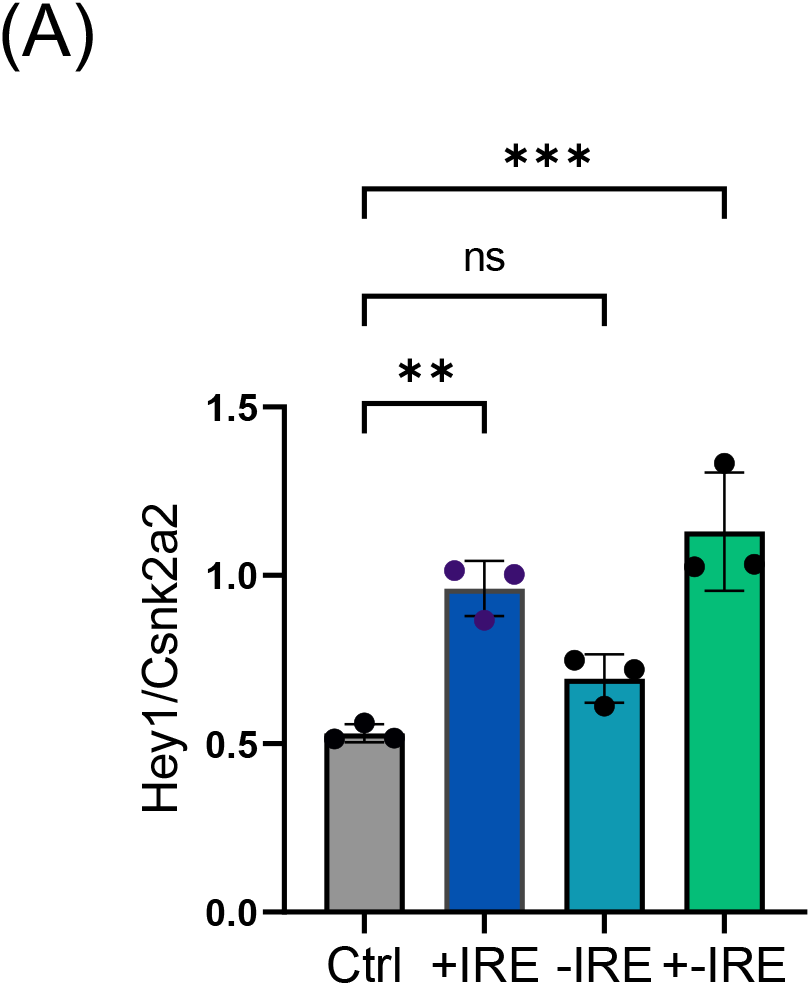
Notch target gene (*Hey1*) expression. **A)** Notch target gene (*Hey1*) expression in Dmt1 KO MEFs with the re-expression of Dmt1 isoforms analyzed by qPCR. One-way ANOVA with Tukey post hoc tests has been performed. Data are represented as mean ± standard deviation. Individual dots represent independent observations. Statistical significance \**p*-value<0.05, \*\**p*-value<0.01, \*\*\**p*-value<0.001 and \*\*\*\**p*-value<0.0001.

## Notes

### Competing Interest Statement

The authors have declared no competing interest.

